## Supplemental Figures for "Regeneration of the larval sea star nervous system by wounding induced respecification to the sox2 lineage"

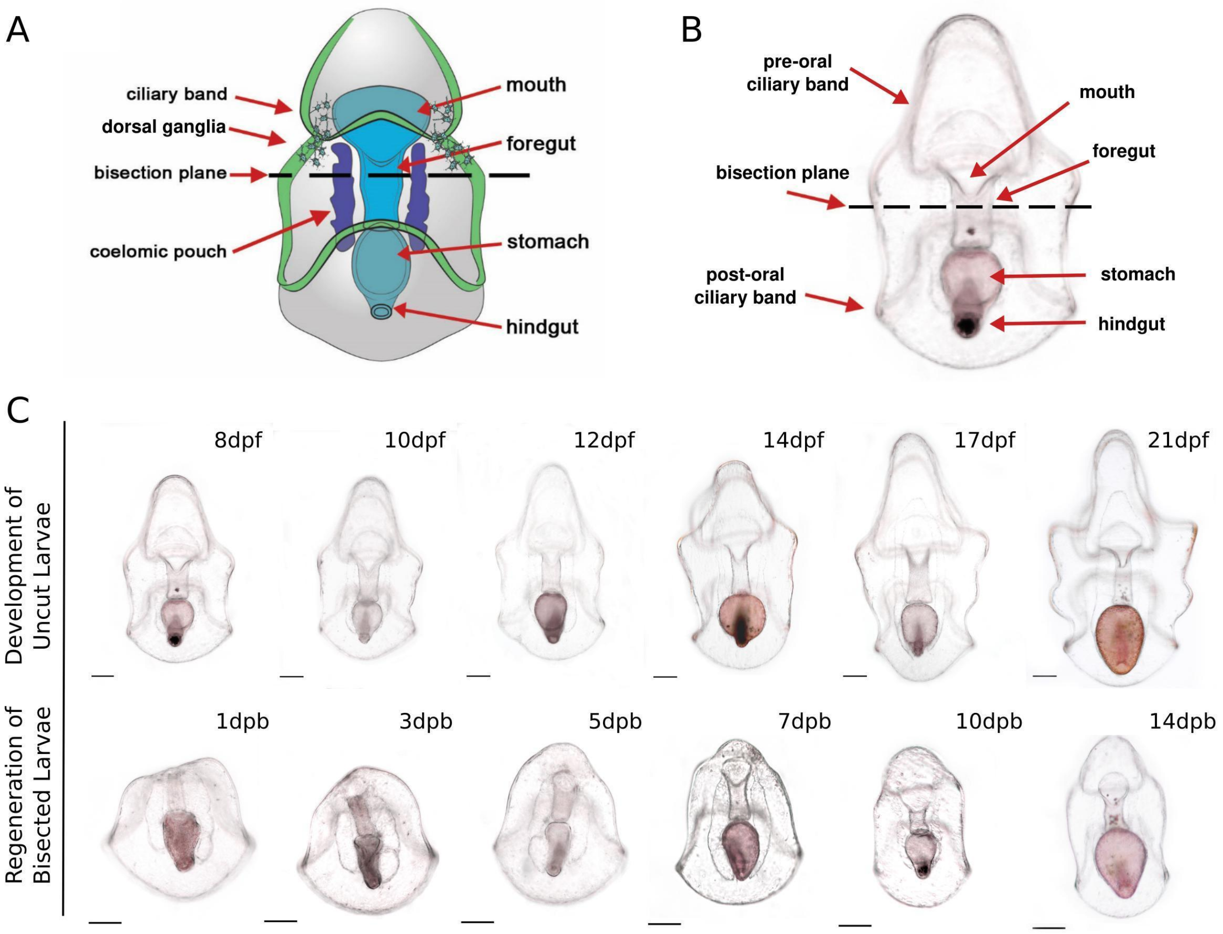


**SP 1. Bright field micrograph showing sea star larvae undergo whole-body regeneration (WBR).**

(A-B) A schematic (A) and bright field micrograph (B) of a 7day post-fertilization (dpf) Patiria miniata sea star larva. Bisection is performed at the midline along the anterior-posterior (AP) body axis beneath the lip of the mouth and through the foregut as indicated by the dotted line. (C) Morphology of posterior sea star regeneration. The panel on top is the morphology of uncut control larvae at corresponding time points. These larvae continue to feed and develop towards a brachiolaria larva. The panel on the bottom is the morphology of regenerating posterior larvae. By 3 days post bisection (dpb), the wound is sealed and the mouth is reformed. In larvae from 3-5 dpb, a pre-oral regeneration leading edge is formed at the anterior. By 7dpb, a primitive anterior structure, the pre-oral ciliary band is formed. The posterior regenerant larvae appear morphologically fully reformed by about 14 days. Scale bar: 100 µm. dpf: day-post-fertilization; dpb: day-post-bisection.

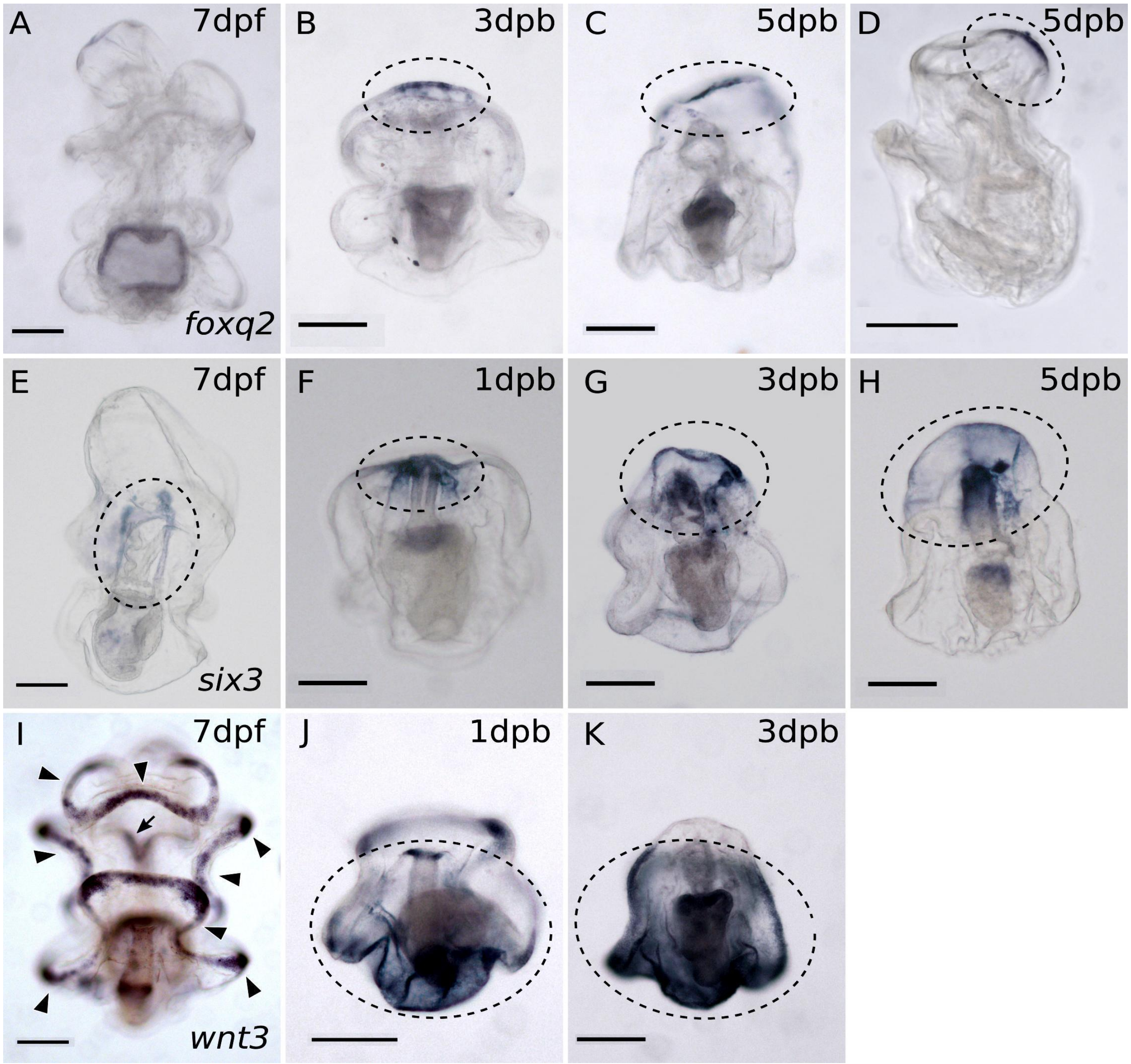


**SP 2. Whole-mount *in situ* hybridization (WMISH) results show the reconstruction of the AP body axis.**

(A-D) WMISH of *foxq2* which is normally expressed in the apical pole domain of the embryo. (A) *Foxq2* expression is undetectable by WMISH in intact larvae (some nonspecific background staining is present in the stomach). Upon decapitation, (B) *foxq2* is first expressed in the anterior-most regeneration leading edge at 3dpb. (C) *Foxq2* expression remains concentrated at the regeneration leading edge by 5dpb. (D) is a lateral view of a regenerating larva. The expression of *foxq2* is detected in the anterior. (E-H) WMISH of *six3* which is normally expressed in the anterior ectoderm of developing embryos. (E) *Six3* is expressed at the bilaterally located coeloms in uncut 7dpf larvae. Upon decapitation, (F) *Six3* expression is detected at the wound ectoderm and the regenerating coeloms in 1 dpb larva. (G-H) *Six3* expression extends to the entire regenerated anterior. (I-K) WMISH of *wnt3* which is normally expressed in the posterior ectoderm of embryos. (I) In intact larvae, *wnt3* is expressed in the ciliary bands, marked by black arrowheads. It is also expressed in the mouth ectoderm indicated by the black arrow. Upon decapitation, (J) *wnt3* is expressed in the posterior domain by 1dpb. (K) The expression extends to the entire *foxq2*-free posterior domain by 3dpb. Gene expression is highlighted in circles with black dashed line. Scale bar: 100 μm. RL, regeneration leading edge; co, coeloms; mo, mouth; CB, ciliary band; D, dorsal; V, ventral.

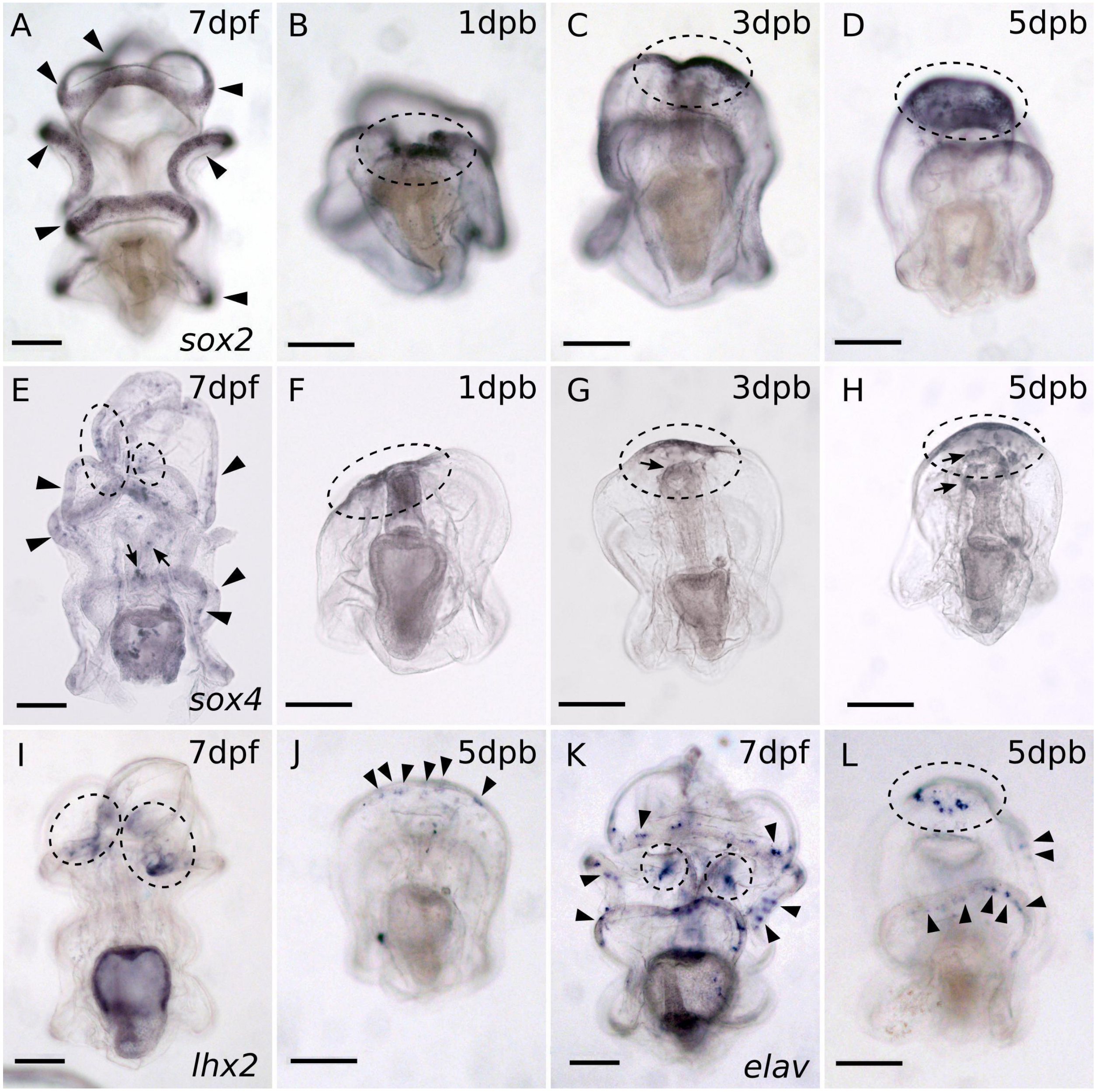


**SP 3. WMISH of embryonic neurogenic pathway genes during regeneration.**

(A-D) WMISH of *sox2.* (A) In intact larvae, *sox2* expression is detected in the ciliary band, indicated by the black arrowheads. (B) Upon bisection, *sox2* is expressed at the wound site and then at the regeneration leading edge as shown in (C-D). (E-H) WMISH of *sox4.* (E) In intact larvae, *sox4* expression is detected in the ciliary band, indicated by the black arrowheads, and in the mouth and foregut, marked by the arrows. *Sox4* is also expressed in the dorsal ganglia, highlighted with black circles. (F) upon bisection, *sox4* expression is detected at the wound site and then in (G-H) at the regeneration leading edge, indicated by circles. In regenerating larvae, *sox4* is also expressed in the oral cells, arrows in (G-H). (I-J) WMISH of *lhx2.* (I) *Lhx2* is expressed at the dorsal ganglia in larvae. (J) In regenerating larvae, *lhx2* expression is first detected at 5 dpb at the regeneration leading edge. (K-L) WMISH of gene *elav.* (K) Post-mitotic neuron marker *elav* is expressed along the ciliary bands (black arrowheads) and the dorsal ganglia (circles). (L) In 5 dpb regenerated larvae, *elav* expression is detected at the regenerated (circle) and remains in the post-oral ciliary band (arrowheads). Scale bar: 100 µm. Dpf: day-post fertilization; dpb: day-post bisection.

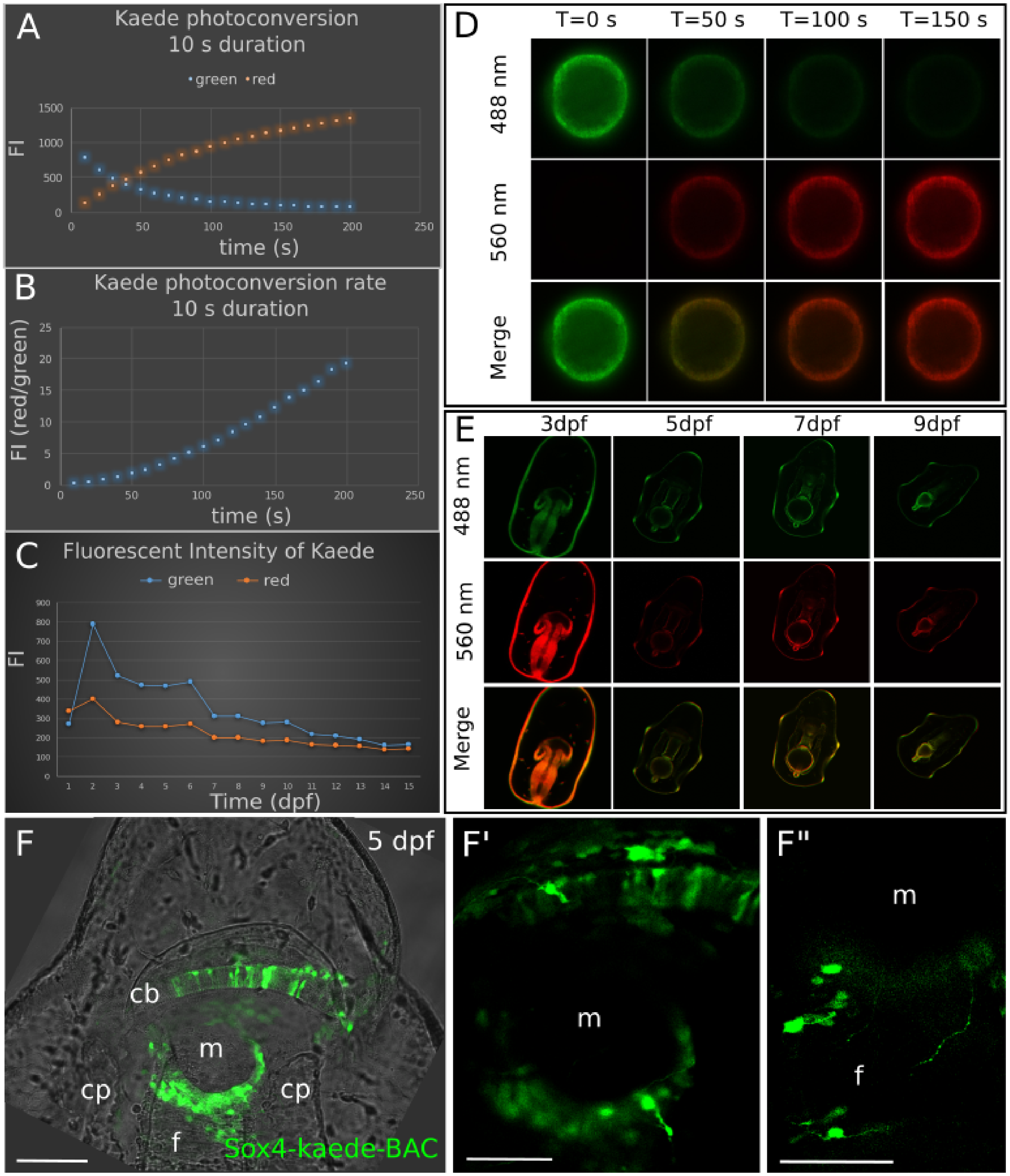


**SP 4. Stability of Kaede protein in P.miniata during development.**
(A-E). Dynamics of photoconversion of Kaede protein. Kaede mRNAs were injected into fertilized eggs where they are endogenously translated by the embryos. Then we photoconverted the Kaede in injected embryos at 24 hpf. The embryos were imaged

every 10s during photoconversion, and then incubated in sea water for at least one week and were imaged again at larval stages. This experiment shows that Kaede proteins unconverted (green) and converted (red) are stably detected above background autofluorescence for at least one week after photoconversion.
(A). shows the fluorescent intensity (FI) of the green and red protein. (B). Photoconversion rate of Kaede protein. This rate is the ratio of the FI (red) over the FI (green). The 560 nm FI gradually increases as the 488 nm FI is progressively reducing with a converging point at ~50s. (C). FI of Kaede protein at different time points. The 488 nm fluorescent intensity rises dramatically at 2 dpf, likely due to the continuous translation of *kaede* mRNA. Red and green proteins in the embryos remain confidently detectable above background in the larvae until at least 7 dpf. Therefore, we conclude that the Kaede proteins can be effectively detected for 7 days. (D) Confocal images of the embryos during the photoconversion. (E). Detection of Kaede in larvae injected with *kaede* mRNA. This shows that Kaede can be stably detected for 7 days. (F-F”). Then we generated a sox4-Kaede BAC to trace the expression of sox4 at larval stages. The expression pattern of sox4-Kaede BAC recapitulates the sox4 mRNA expression data (SP. Fig.3, E). Some of the sox4+ cells have clear neural phenotypes. Scale bar in (F): 100 μm; (F’-F”): 50 μm. Cb, ciliary band; cp, coelomic pouch; f, foregut; m, mouth; dpf, day post fertilization.

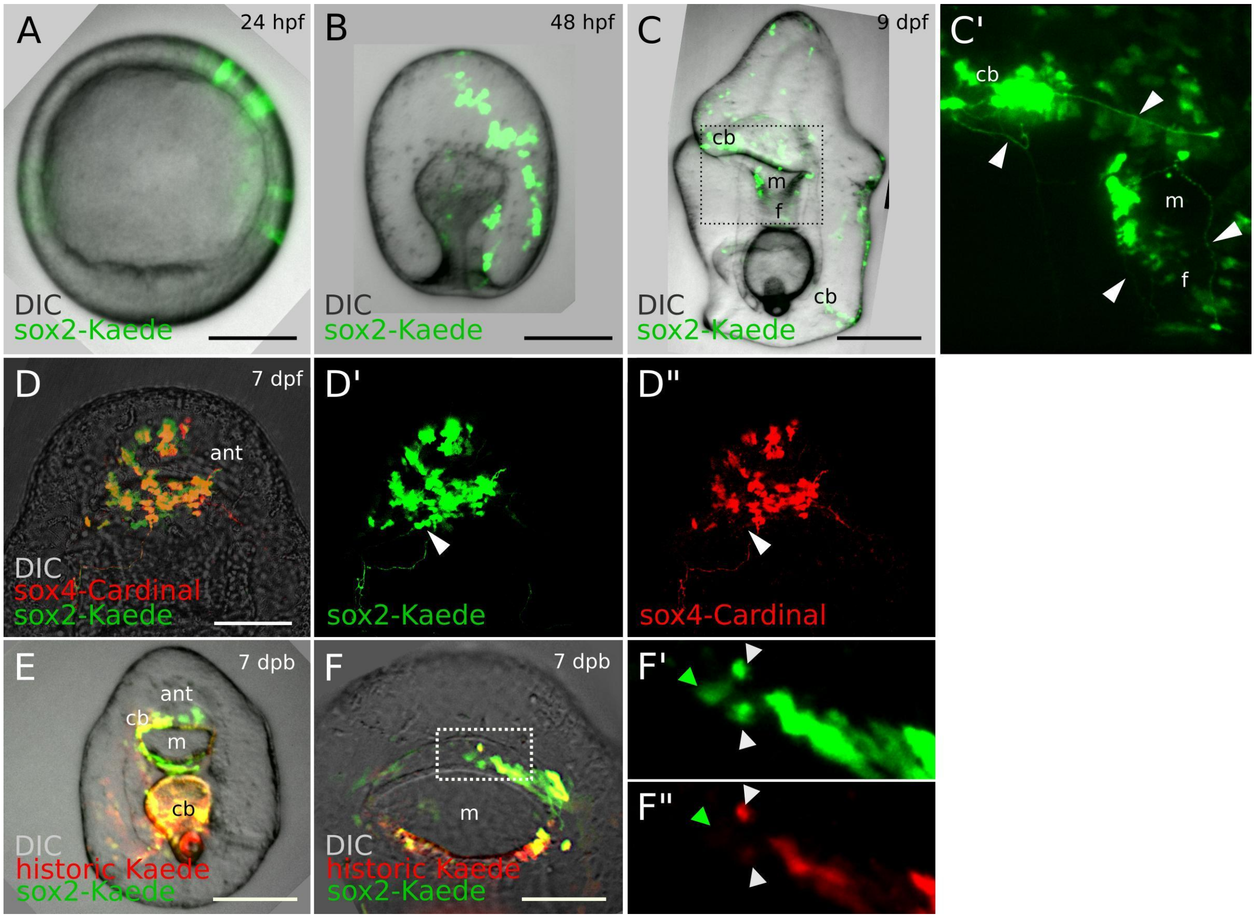


**SP 5. Expression pattern of sox2-Kaede BAC.**

(A-B). Sox2 transgenic embryos. (C-C'). In the larval stage sox2+ cells are found in the ciliary band, mouth, and foregut. Some cells have clear long, axonal projections characteristic of neural morphology (arrowheads). The boxed area in (C) amplifies in C'. (D-D"). Normal transgenic larvae coexpressed two BACs: sox2:Kaede and sox4:Cardinal. This shows that the majority of sox2+ cell lineage are neurogenic and give rise to ectodermal sox4+ cells. Some sox2+ cells (white arrowheads) do not express sox4 BAC. (E-F"). In 7dpb regenerating larva sox2+ cells reused (yellow) and de novo (green, F', F") located in the restored mouth, ciliary band. The boxed area in (F) amplifies in F', F". Scale bars in (A, B): 25 µm, in (C, D, E, F): 50 µm. Ant, anterior; cb, ciliary band; f, foregut; m, mouth.

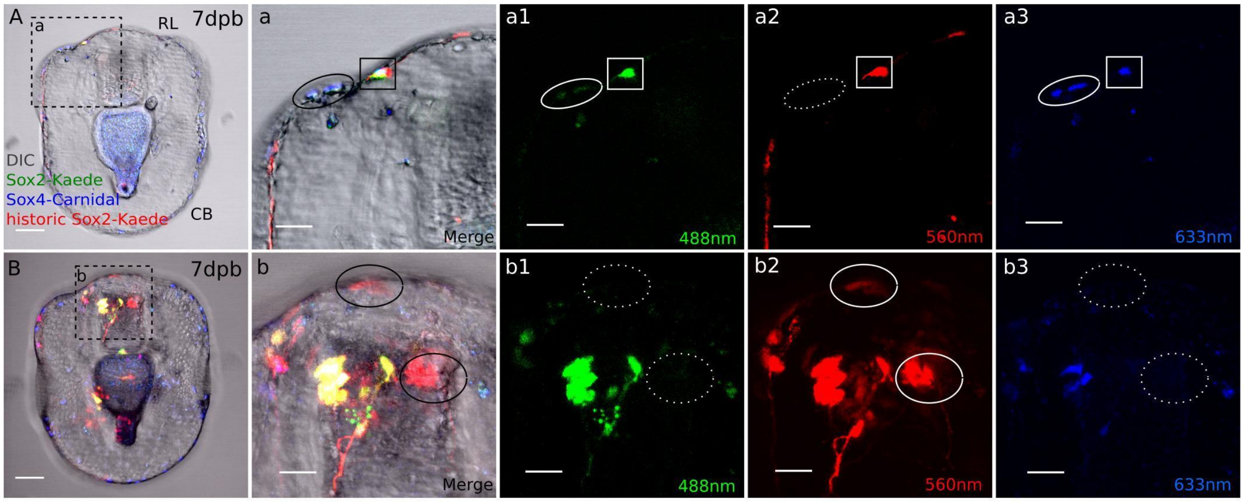


**SP 6. Sox2+ cell lineage forms the regenerated nervous system.**

(A). *De novo* sox2 cells form neurons through the sox4-mediated pathway in the regenerating anterior. (a) Amplification of boxed area in (A). There are two neurons at the lateral side of the regenerating anterior (black solid circle) derived from the newly specified sox2+/sox4+ cells. They contain (a1) regenerative green sox2-Kaede and (a3) blue sox4-Cardinal (white solid circle), (a2) but do not have historic sox2 expression (dotted white circle). Other than the *de novo* sox2+ cells, the existing sox2+ lineages also contribute to the regenerated neurons at the anterior marked by the black solid square in (a). (a1-a3). This neuron contains all three colors (white solid box), suggesting it is derived from the existing sox2+ cell lineage through sox4-mediated pathway. (B). Historic sox2+ cells present neural morphology. (b). There are red only, historic sox2+ neural cells at the regeneration leading edge and the oral domain (black solid circles). These cells are likely to be differentiated cells or cells taken other fates in regeneration, thus no longer express sox2. (b2). They contain the historic Kaede marker, (b1) but do not express new, green Kaede or (b3) Cardinal (white dotted circle). Scale bar in (A-B): 50 µm. RL, regeneration leading edge; CB, ciliary band.

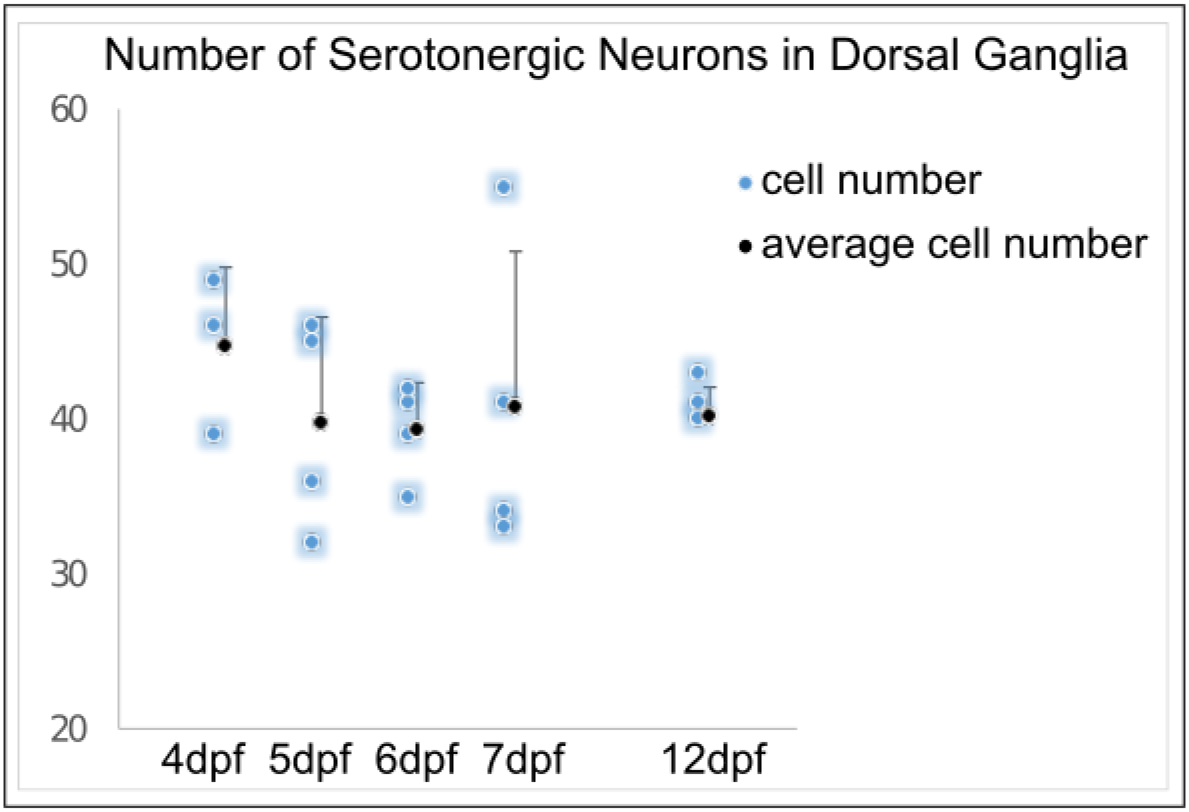


**SP 7. Number of serotonergic neurons in larvae**

Quantification of serotonergic neurons in larvae shows a stable number of serotonergic neurons over time. Serotonergic neurons were counted manually in Z-stack images of 5-HT immunostaining of larvae at different time points using Fiji image processing software.

**SP Table. List of used antibodies and BACs.**

Table 1. Antibodies used in this study

| Antibodies | Source | Host species | Dilution |
|---|---|---|---|
| Primary antibodies | | | |
| Serotonin | Sigma (S5545) | rabbit | 1:250 |
| 1E11 | Nakajima et al., 2004 | mouse | 1:5 |
| Secondary antibodies | | | |
| Goat anti-mouse Cy3 | Jackson Immunoresearch (115-165-146) | goat | 1:2000 |
| Goat anti-rabbit Cy3 | Jackson Immunoresearch (115-165-144) | goat | 1:2000 |
| Goat anti-rabbit Alexa Fluor 488 | Invitrogen (A11008) | goat | 1:1000 |

Table 2. BACs used in this study

| Gene of interest | Fluorescent reporter | Source |
|---|---|---|
| Sox4 | GFP | echinobase.org |
| | Kaede (Kaede-N1, Addgene #54726) | |
| | mCardinal | |
| Sox2 | Kaede (Kaede-N1, Addgene # 54726) | |
